# Coevolution with toxic prey produces functional trade-offs in sodium channels of predatory snakes

**DOI:** 10.1101/2023.12.08.570760

**Authors:** Robert E. del Carlo, Jessica S. Reimche, Haley A. Moniz, Michael T.J. Hague, Shailesh R. Agarwal, Edmund D. Brodie, Edmund D. Brodie, Normand Leblanc, Chris R. Feldman

## Abstract

Seemingly unrelated traits often share the same underlying molecular mechanisms, potentially generating a pleiotropic relationship whereby selection shaping one trait can simultaneously compromise another. While such functional trade-offs are expected to influence evolutionary outcomes, their actual relevance in nature is masked by obscure links between genotype, phenotype, and fitness. Here, we describe functional trade-offs that likely govern a key adaptation and coevolutionary dynamics in a predator-prey system. Several garter snake (*Thamnophis* spp.) populations have evolved resistance to tetrodotoxin (TTX), a potent chemical defense in their prey, toxic newts (*Taricha* spp.). Snakes achieve TTX resistance through mutations occurring at toxin-binding sites in the pore of snake skeletal muscle voltage-gated sodium channels (Na_V_1.4). We hypothesized that these mutations impair basic Na_V_ functions, producing molecular trade-offs that should ultimately scale up to compromised organismal performance. We investigate biophysical costs in two snake species with unique and independently evolved mutations that confer TTX resistance. We show electrophysiological evidence that skeletal muscle sodium channels encoded by toxin-resistant alleles are functionally compromised. Furthermore, skeletal muscles from snakes with resistance genotypes exhibit reduced mechanical performance. Lastly, modeling the molecular stability of these sodium channel variants partially explains the electrophysiological and muscle impairments. Ultimately, adaptive genetic changes favoring toxin resistance appear to negatively impact sodium channel function, skeletal muscle strength, and organismal performance. These functional trade-offs at the cellular and organ levels appear to underpin locomotor deficits observed in resistant snakes and may explain variation in the population-level success of toxin-resistant alleles across the landscape, ultimately shaping the trajectory of snake-newt coevolution.

## Introduction

Organismal performance is the integration of many intervening phenotypic steps that translate genetic variation into fitness^1^. The genetic changes required to adapt to novel selective pressures must therefore integrate with preexisting molecular mechanisms, cellular processes, and physiological functions that cooperate to maintain organismal performance^1^. Consequently, when selection acts on multiple phenotypes controlled by a single gene, mutations capable of producing phenotypes that overcome one selective demand must also largely preserve the encoded protein’s structure so as to maintain all other molecular functions. When very few, if any, substitutions can accommodate this constraint, mutating a protein that underlies multiple key traits can produce phenotypic trade-offs, whereby adaptive change in one trait jeopardizes another (antagonistic pleiotropy) ^2–5^. Biophysical constraints on protein structure and function can therefore impose limits on adaptive evolution ^2,5–8^. As such, decomposing organismal fitness into its constituent molecular phenomena is necessary to understand how phenotypic trade-offs might constrain adaptive evolution, ultimately determining the degree to which evolutionary trajectories are predictable across lineages^9^.

Here, we demonstrate trade-offs at hierarchical levels of biological organization in a classic coevolutionary system involving toxic newt prey (*Taricha* spp.) and predatory garter snakes (*Thamnophis* spp.). Newts employ tetrodotoxin (TTX) as an anti-predator defense^10–12^, yet some populations of garter snakes have evolved resistance to this potent neurotoxin^13–16^. Resistance is conferred by mutations directly to the TTX-binding site on voltage-gated sodium channels (proteins: Na_V_; genes: *SCNnA*)^13,17–21^. While these structural changes produce effective resistance to TTX, they occur in highly conserved regions of the protein that are essential to channel function^22^. Therefore, we hypothesized that the highly conserved structure-function relationship of this critical protein constrains adaptive evolution in this system, whereby mutations that increase TTX resistance are deleterious to channel function and, by extension, physiological and whole-animal performance^23–25^.

Sodium channels are transmembrane proteins expressed in nerves and muscles that carry vital sensory and motor impulses^26–28^. The 24 transmembrane passes of these proteins form four voltage-sensing domains (see Figure 2a) that dilate a central, sodium-selective pore through which sodium ions enter the cell, carrying a positive charge that depolarizes the membrane^29^. TTX occludes the Na_V_ outer pore, thereby arresting nervous and mechanical activity, leading to paralysis and death^30,31^. This poison’s chemical structure is so compatible with the topology of the nearly universally conserved Na_V_ pore that just nanomolar concentrations of TTX are lethal to nearly all animals^30,32^. Consequently, newts exploit TTX as a broad-spectrum defense to evade predation, secreting the poison from their skin when attacked^33,34^, and paralyzing or killing virtually all potential predators^11^.

Despite this, TTX resistance has evolved repeatedly in nature, generally through target-site insensitivity conferred by mutations directly to the outer pore of the sodium channel^19,23,35–38^. However, mutations to Na_V_ can have dire phenotypic consequences because many Na_V_ functions are tied directly to protein structure, including selectivity to sodium, the rate of sodium influx by electrodiffusion, and sensitivity to changes in membrane potential ^29,39–41^. As such, sodium channel mutations are particularly hazardous, often resulting in serious congenital disease or stillborn progeny^42^. Dietary TTX has nevertheless driven the evolution of snakes carrying very effective Na_V_ mutations, though functional variation is limited to just a few pore sites (Fig. 1a,d,g) ^13,23,32^. Thus, the tight relationship between molecular structure and sodium channel function is thought to constrain NaV evolution^32^, and underlie the remarkable convergent molecular evolution seen across TTX-resistant taxa^23,25^.

**Figure 1.**
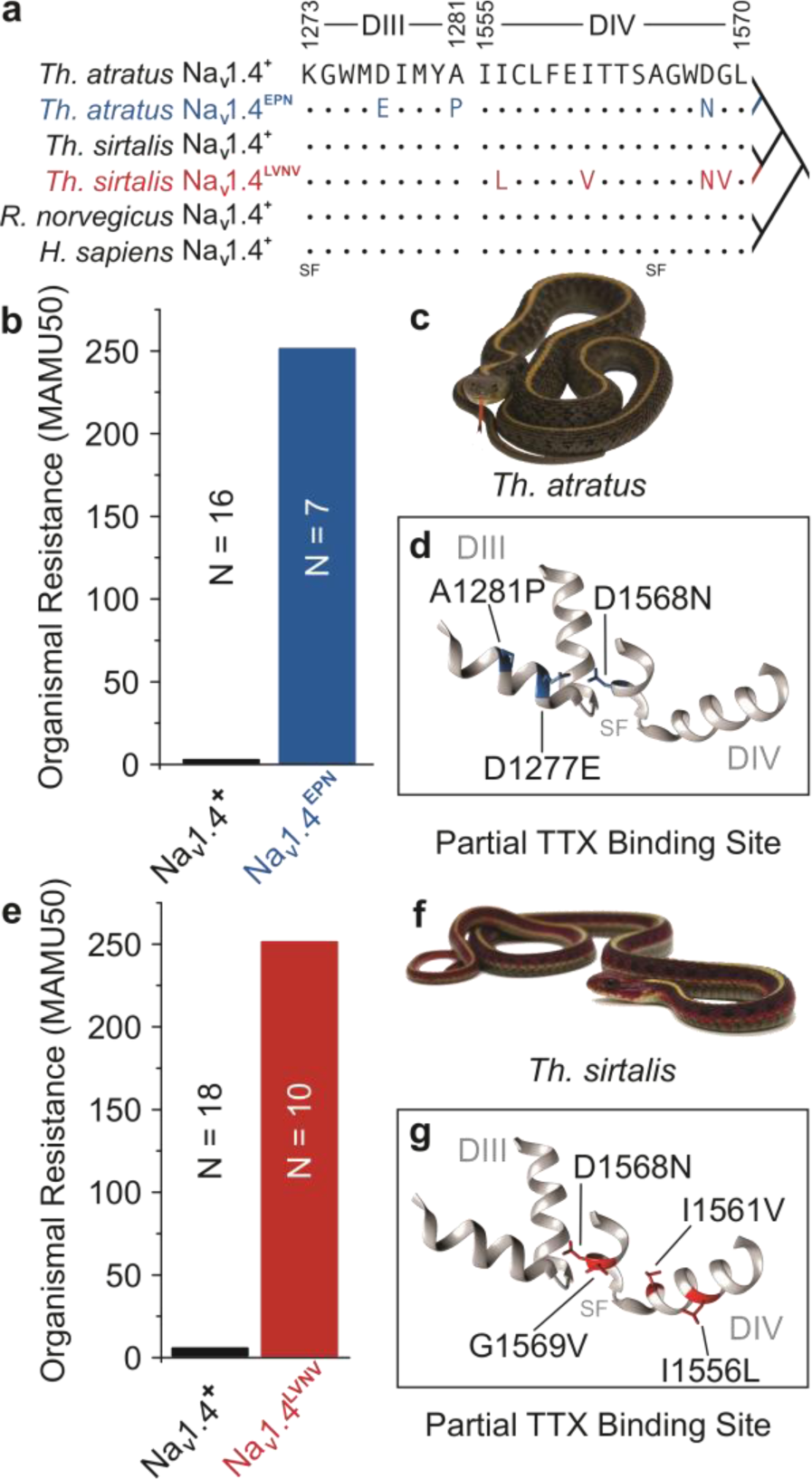
Pore domain mutations found in extremely TTX-resistant snakes. **a**, The pore-forming domain of Na_V_1.4 is highly conserved across disparate phyla, although some populations of *Thamnophis* spp. exhibit mutations in pore domains III and IV very closely opposed to the selectivity filter (SF) (GenBank: *Rattus norvegicus* NM_013178.1 and *Homo sapiens* NP_000325.4)^23^. **b,e**, Snakes in both species carrying pore mutations are extremely resistant to TTX relative to their conserved counterparts. Resistance is measured here as the median dose required to reduce snake locomotor activity to 50%, scaled relative to the dose at which a mouse of equal mass would succumb (50% Mass-Adjusted Mouse Unit, MAMU50). Whereas a dose of 1 MAMU subdues mammals, mutant populations of *Th. atratus* (**c**) and *Th. sirtalis* (**f**) are 250X more resistant to TTX. We designate snakes carrying conserved pore sequences as TTX-sensitive despite being somewhat more resistant than mammals (*Th. atratus*: 1.35 MAMU50 and *Th. sirtalis*: 4.16 MAMU50). Medians are presented as neither resistant population shows variation in this metric and the variation in sensitive populations is imperceptible at the scale presented. Pore domain mutations deform the TTX binding site (**d, g**), thereby conferring resistance. However, the high degree of conservation at these substitution sites implies a high degree of functional significance that likely underlies a phenotypic trade-off in this system.

The requirement that voltage-gated sodium channels must maintain essential electrophysiological performance while simultaneously preventing TTX blockade therefore imposes competing selective demands on a single locus. Thus, resistant sodium channels are expected to suffer from antagonistic pleiotropy^23,25,43^. Indeed, previous correlational work^15,24,44^ suggests that some populations of TTX-resistant snakes suffer locomotor deficits. However, the etiology of whole-animal performance costs remains obscure and has only been considered for a single group of TTX-resistant snakes^24,43^. Work is needed to directly test the impact of naturally occurring Na_V_ mutations, from the biophysical level to higher order physiology. To determine whether antagonistic pleiotropy creates functional trade-offs that may influence evolutionary outcomes, we need to understand how resistance-conferring mutations disrupt the sodium channel structure-function relationship, resulting in a cascade of performance deficits from molecules to phenotypic trade-offs at the organismal level^24,44^.

To identify the biophysical mechanism of higher order trade-offs, the molecular effects of TTX-resistant channels must be quantified and then connected to relevant physiological systems (*i.e.* skeletal muscle) to develop a framework that explains trade-offs at the whole-animal level^1,24,43,45^. Here, we assessed the biophysical costs of resistance-conferring mutations to the skeletal muscle voltage-gated sodium channel (Na_V_1.4), primarily finding mutant channels produce less bioelectricity (reduced unitary conductance) and are therefore less able to excite skeletal muscle compared to their TTX-sensitive counterpart. We then investigated mechanical deficits that scale up from effects seen at the molecular level, revealing reduced force output and slowed mechanical activity of skeletal muscle in toxin-resistant snakes. Lastly, we examine how these structural mutations alter the interaction energy dynamics of functional residues in the sodium channel, offering insight into the molecular mechanism of this phenotypic trade-off. The evidence of this trade-off is reproducible across two independently evolved resistant snake species with unique resistance alleles and may underlie the heterogeneous landscape-level distribution of toxin resistance^16,20,46,47^. We conclude that performance trade-offs at the molecular level are the proximate source of higher-order limitations that give rise to constraints on natural selection, and in so doing, comprise a straightforward map from gene sequence to evolutionary consequence.

## Results

### Resistant channels produce less current than their toxin-sensitive counterpart

We tested the hypothesis that mutations in the highly conserved sequence of the sodium channel pore (Fig. 1a, d, g; Fig. 2a) underlie organ- and higher-level performance deficits. We characterized two suites of mutations in skeletal muscle voltage-gated sodium channels (Na_V_1.4) associated with the greatest levels of organismal TTX resistance seen in two species of *Thamnophis*^23^. We assessed the electrophysiology of the conserved, TTX-sensitive channel (Na_V_1.4^+^, *Rattus norvegicus*) and two TTX-resistant channel mutants: Na_V_1.4^EPN^ (D1241E, A1245P, and D1532N) and Na_V_1.4^LVNV^ (I1520L, I1525V, D1532N, and G1533V) (Fig. 2b). Mutations at these positions on the *Rattus* channel backbone recreate pore loop (p-loop) substitutions found at analogous sites in wild, TTX-resistant snake Na_V_1.4 Domains III and IV of *Th. atratus* (D1277E, A1281P, and D1568N) and Domain IV of *Th. sirtalis* (I1556L, I1561V, D1568N, and G1569V) (Fig. 1a). As expected, we found that these snake mutations confer strong TTX resistance to the rat channel relative to the conserved protein (Fig. 2c). In search of a possible source of trade-offs, we began by assessing the activation and inactivation parameters of sensitive and resistant channels.

**Figure 2.**
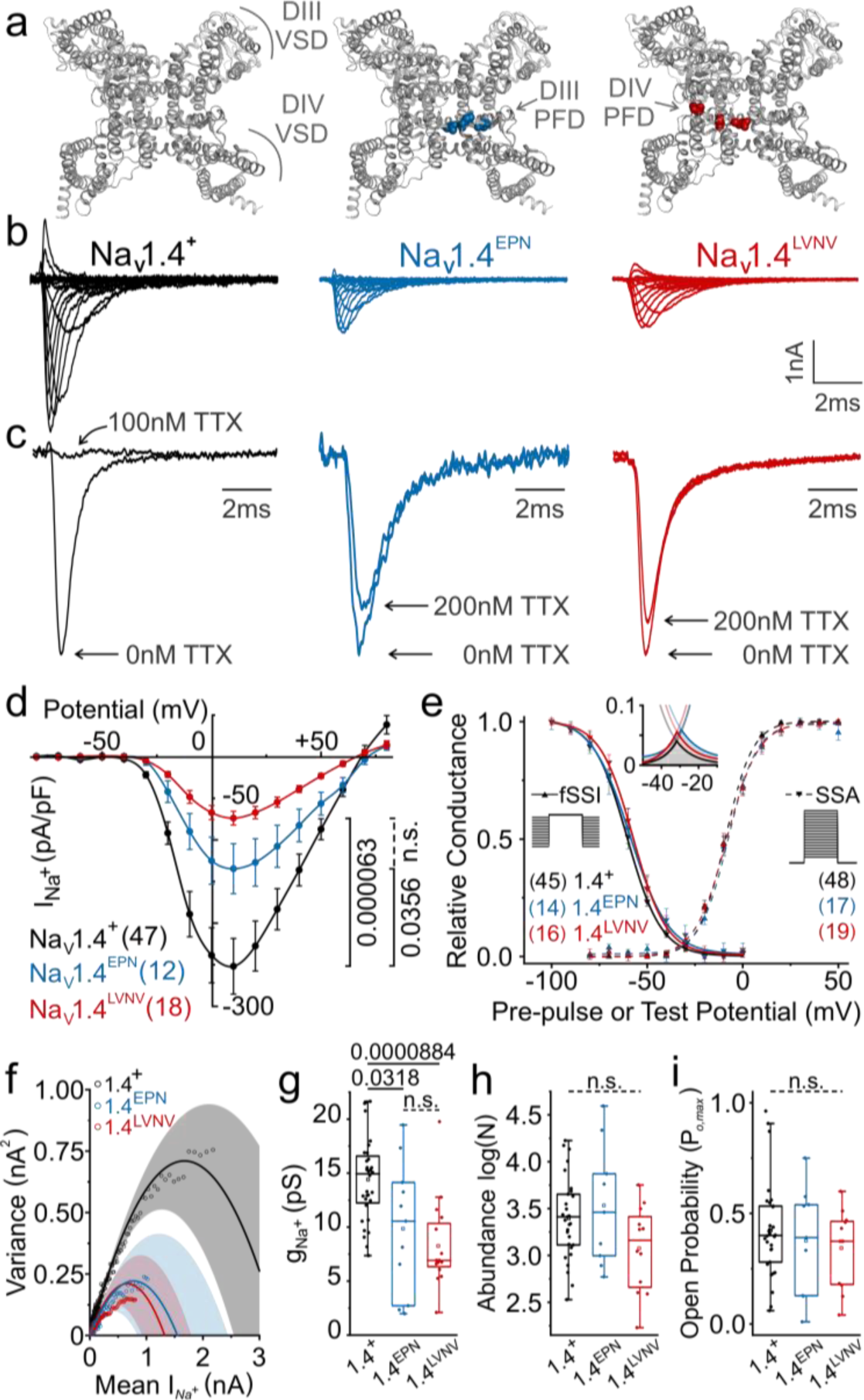
Sodium channel unitary conductance is reduced in two independently evolved pore domain mutants that confer TTX resistance. **a**, Tertiary structures of skeletal muscle voltage-gated sodium channels from the extracellular face (*Homo sapiens* structure, PDB: 6agf) showing the TTX-sensitive Na_V_1.4^+^ (black) as well as the TTX-resistant *Th. atratus* Na_V_1.4^EPN^ (blue) and *Th. sirtalis* Na_V_1.4^LVNV^ (red) mutations. Movement of the voltage-sensing domains (VSDs) dilates the sodium-permeant central pore by tugging on the pore-forming domains (PFDs), the surface of which forms the binding site for TTX. **b**, Representative families of macroscopic sodium currents recorded from *Rattus* clones carrying identical mutations. **c**, Mutagenized rat channels are hardly affected by relatively large doses of TTX. **d**, Current magnitude among TTX-resistant sodium channels is reduced without any differences in (**e**) voltage-dependence of activation, inactivation or peak window current (**e** inset). **f**, Current-variance relationships demonstrate deficits in unitary current (cord conductance, **g**) among TTX-resistant sodium channels without any concomitant differences in sodium channel number (*N*, **h**) or peak-open probability (P_o,max_, **i**). All p-values presented are calculated by Dunn’s post-hoc pairwise comparison test after Kruskal-Wallis nonparametric ANOVA. Significantly different pairwise comparisons denoted by solid bars and their associated p-value; nonsignificant results (n.s.) denoted by dashed lines.

Sodium channels activate in response to depolarizing membranes. Mutations can change the flexibility of the protein, thereby potentially dampening or facilitating the channel’s conformational changes in response to depolarization. Muscle weakness could arise from mutations that cause the sodium channel to be less responsive to depolarization (“hypoexcitability”). The resting membrane potential of skeletal muscle is very negative, varying across phyla from -70mV to nearly as low as -90mV^48–50^. At such negative resting potentials, healthy sodium channels should remain closed, but ready to open^51^. When an organism intends to contract a voluntary skeletal muscle, peripheral nerves release acetylcholine onto the neuromuscular junction, depolarizing the postsynaptic membrane through the action of ionotropic Nicotinic Acetylcholine Receptors (nAChR)^52^. The brief currents raise the local transmembrane voltage. For any sodium channels near enough to be under the electrical influence of this depolarization, the positively-charged residues in the voltage-sensing domains of Na_V_ are repelled extracellularly by this more positive (less negative) internal potential. The conformational change in the voltage-sensing domains applies torsion to the pore-forming domains, dragging them radially outward, thus dilating the channel enough to conduct hydrated sodium ions, producing a current that propagates the depolarizing wavefront^53–55^. The voltage at which a population of sodium channels opens in near unison defines the “threshold” of an action potential. Channels exhibiting a higher threshold (lower sensitivity to depolarization) will result in muscles that require a greater stimulus to produce a contraction. Thus, the voltage sensitivity of sodium channel activation is a reasonable target for the investigation of trade-offs in toxin resistant snakes.

In whole-cell patch clamp electrophysiology experiments where transmembrane voltage is controlled by an external amplifier, the population of channels in the membrane is held at a constant voltage, conditioning the proteins into nearly uniform molecular configurations. At potentials near or more negative than rest, the voltage-sensing domains are pulled intracellularly, cinching the pore-forming domains, preventing channel patency^56–58^. As test potentials are increasingly depolarized (*i.e.* sequentially more positive), greater and greater proportions of the sodium channel population will respond to the activating stimulus, resulting in macroscopic sodium currents that are larger in magnitude. The development of sodium currents from a population of channels conditioned into an equilibrium of nearly uniform conformational states is referred to as “steady-state activation.”

Any mutation near the pore or voltage-sensing domains could feasibly alter the rigidity or flexibility of this orchestrated molecular reconfiguration, thereby requiring greater or lesser depolarizing potentials to achieve the same magnitude of sodium current. However, we found that the voltage dependence of steady-state activation is identical across all sodium channel variants observed in this study (Fig. 2e). That is to say, across a range of membrane potentials from -80mV to +80mV, equivalent proportions of both sensitive and resistant sodium channel populations respond to the same depolarizing stimulus. Therefore, voltage sensitivity of activation does not explain higher order locomotor deficits.

In addition to activating in response to depolarization, healthy sodium channels inactivate in time- and voltage-dependent manners^59^. Under normal circumstances, this is achieved by a hydrophobic intracellular loop that becomes mobile once the channel is activated, contorting to occlude the channel pore from the intracellular face, preventing further passage of sodium despite voltage-sensing domains remaining in the activated conformation^60^. The total macroscopic current decays within a few milliseconds as inactivation stochastically sets in for all sodium channels throughout the membrane, known as “fast inactivation”^61,62^. Slow inactivation takes place over much longer time scales and provides a less convincing etiology for trade-offs in toxin-resistant snakes^61^. Once inactivated, sodium channels are very unlikely to return to a conducting state until the membrane potential returns to near or below rest, resetting the voltage-sensing domains and removing the hydrophobic inactivation gate. If a mutation were to cause premature inactivation, perhaps by increasing the mobility of the inactivation gate, then less overall current would pass through the membrane, thereby limiting the depolarizing power of the sodium channel and hampering the excitability and contractility of skeletal muscle.

TTX resistance mutations in snakes are found on the extracellular face of the channel pore, separated from the inactivation gate on the intracellular face^63^. Therefore, pore-domain mutations are not known to affect the voltage-dependence of fast inactivation, which is consistent with our findings here (Fig. 2e)^43,64,65^. However, holding voltage constant, we do see that one resistant variant inactivates more quickly than others (Supp. Fig. 1g,h). While the triple-point mutant (*R.n.*Na_V_1.4^EPN^) exactly follows the inactivation rate of the ancestral TTX-sensitive channel (*R.n.*Na_V_1.4^+^), the quadruple-point mutant (*R.n.*Na_V_1.4^LVNV^) enters steady-state inactivation significantly faster than the TTX-sensitive channel. This premature onset of fast inactivation in one toxin-resistant channel may underlie higher-order physiological deficits unique to *Thamnophis sirtalis* (described later). Assessing how quickly inactivation can be removed in these mutant channels, we find that neither the rate (Supp. Fig. 2) nor the voltage dependence of recovery from inactivation (Supp. Fig. 2) are affected by these pore-domain mutations. Thus, these distinct resistance-conferring p-loop mutations appear to exert unique effects on some channel functions while entirely sparing others. However, with identical steady-state activation and faulty inactivation in only one of the two mutants, these findings alone are insufficient to explain all locomotor deficits in both *Thamnophis atratus* and *Thamnophis sirtalis*.

Without evidence supporting a kinetic explanation for locomotor deficits (Supp. Fig. 1a-d), then the magnitude of sodium currents, and thus the excitability and contractility of skeletal muscle, must be determined by other fundamental properties of Na_V_. Across the current-voltage relationship, TTX-resistant channels produce far less macroscopic current than their toxin-sensitive, conserved counterpart (Fig. 2d, 30% less in *R.n.*Na_V_1.4^EPN^ and 50% less in *R.n.*Na_V_1.4^LVNV^). However, this global, macroscopic current can be decomposed into three variables: how many channels are active in the membrane (N), the maximum statistical likelihood of finding any of the N channels open (P_o,max_), and how much sodium current each individual channel conducts when it is actually open (unitary conductance, or g_Na_)^66^. The most expedient electrophysiological technique to measure these variables is Non-Stationary Noise Analysis (NSNA, Fig. 2f)^66^. Assuming all sodium channels encoded by the same allele perform roughly identically, then any variation in macroscopic current not otherwise attributable to thermal, electromagnetic, or vibrational noise, should be quantal: one open sodium channel always produces the same amount of current. Therefore, by taking an average of the inactivating phase of an ensemble of 1000 sodium currents evoked under identical conditions, the statistical variance of the inherent noise around the mean is proportional to the unitary current of the sodium channel. If activation and inactivation determine at what voltages sodium channels open and for how long they remain open, then the unitary current determines the amount of sodium that can flow per unit time and likely underlies the differences in macroscopic currents observed here between ancestral and mutant channels (Fig. 2d).

Through NSNA, we found median single channel conductance, g_Na_, (Fig. 2g) to be significantly reduced in toxin-resistant mutants (29.3% in *R.n.*Na_V_1.4^EPN^, *p* = 0.0318; 53.6% in *R.n.*Na_V_1.4^LVNV^, *p* = 8.84×10^-5^) without significant changes in channel number, N, (Fig. 2h, Supp. Fig. 1f) or peak open probability, P_o,max_ (Fig. 2i, Supp. Fig. 1.e). Consequently, these mutations appear to reduce the total ionic current by reducing unitary conductance, g_Na_, as opposed to reduced channel expression or activity. Furthermore, the degree of biophysical dysfunction reported here is very likely to have consequences at higher levels of organization capable of producing the muscular and locomotor deficits observed in previous work^24^. The electrophysiological data presented here demonstrate that TTX resistance achieved by target-site insensitivity in Na_V_1.4 is inherently costly by way of reduced sodium channel conductance.

### Resistant muscles develop less force than their toxin-sensitive counterparts

Sodium channels propagate the action potential wavefront across the sarcolemma and into the T-tubules, activating voltage-sensing proteins (Dihydropyridine Receptors, DHPR) that physically couple to and activate Ryanodine Receptors (RyR_1_) in the sarcoplasmic reticulum^52,67,68^. The opening of RyR_1_ releases sarcoplasmic calcium stores into the cytosol where Ca^2+^ can interact with troponin, thereby engaging the contractile machinery^69^. The greatly reduced conductance of TTX-resistant channels likely limits their ability to depolarize the sarcolemma. We therefore hypothesized that sodium channels generating less conductance will fail to maximally engage this cascade of contractile mechanisms, resulting in weaker and slower muscles.

We contrasted the physiological performance of skeletal muscle from individual *Th. atratus* and *Th. sirtalis* carrying ancestral TTX-sensitive pore sequences (*Th.*spp.Na_V_1.4^+^) to their three- and four-point mutant counterparts (TTX-resistant *Th.a.*Na_V_1.4^EPN^ and *Th.s.*Na_V_1.4^LVNV^). We first established the concentration-dependent effect of TTX on isolated skeletal muscle twitch force (Fig 3a,e, Supp. Fig. 4a,f). Muscles from both *Th. atratus* and *Th. sirtalis* carrying the ancestral pore-domain sequences were paralyzed by low concentrations of TTX (IC_50_ ∼ 90-150 nM), while those carrying p-loop mutations withstood orders of magnitude greater TTX concentrations (as much as 100-fold) with many individual muscles continuing to contract at concentrations exceeding 70μM. We observed a strong correlation between TTX resistance in skeletal muscle tissue and resistance in the live animal (adjusted R^2^ = 0.5427, *p* = 1.4×10^-9^, df = 46).

**Figure 3.**
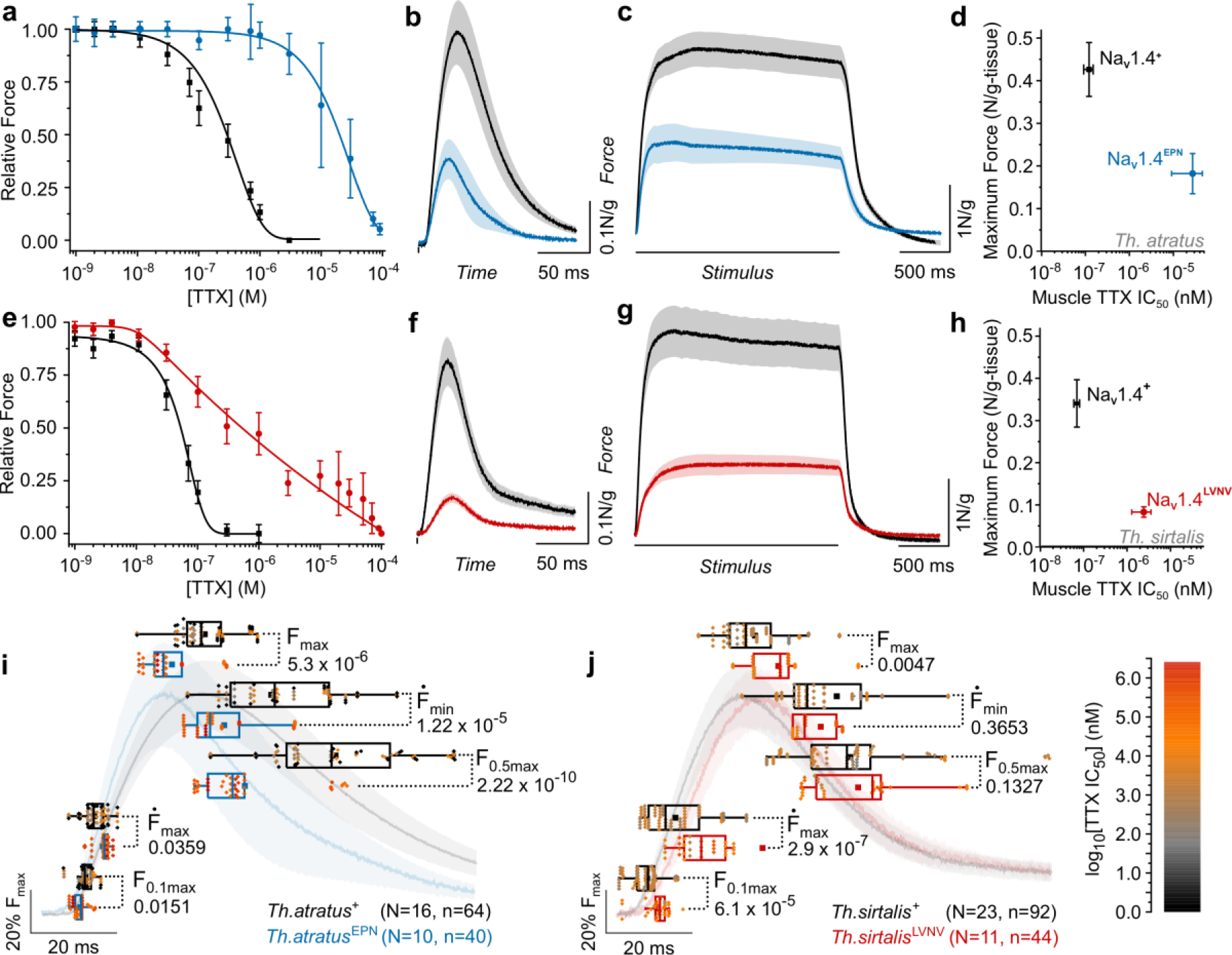
Muscle performance is reduced in two snake species carrying independently evolved TTX-resistance mutations in Na_V_1.4. **a**,**e**, Mean effect of tetrodotoxin concentration [TTX] on snake skeletal muscle transient force (±sem). Data are grouped by species and genotype: *Th. atratus* (**a**) carrying TTX-sensitive Na_V_1.4^+^ (black, N=16, n=64) and TTX-resistant Na_V_1.4^EPN^ (blue, N=10, n=40) and *Th. sirtalis* (**e**) carrying TTX-sensitive Na_V_1.4^+^ (black, N=23, n=92) and TTX-resistant Na_V_1.4^LVNV^ (red, N=11, n=44). **b**,**f**, Transient muscular contractions (mean±sem) with electric field stimulus onset and duration indicated by a vertical black line (grouped as above). **c**,**g**, Tetanic muscular contractions (mean±sem) with a 2 second stimulus indicated by the black line (**c**, *Th.atratus* Na_V_1.4**^+^**N=15, Na_V_1.4^EPN^ N=10; **g**, *Th. sirtalis* Na_V_1.4**^+^**N=17, Na_V_1.4^LVNV^ N=11). **d**,**h**, In both snake species, skeletal muscles carrying ancestral sodium channel pore sequences exhibit greater force while mutant muscles display orders of magnitude greater TTX resistance but weaker force. This is the clearest evidence of a trade-off in populations that have coevolved with tetrodotoxic newts. X-error caps exaggerated for visibility. **i**,**j**, The temporal progression of transient contractions reveals variable timing between genotypes within species. Relevant metrics of contraction chronology include time to 10% contraction (F_0.1max_), time to peak first derivative (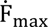), time to 50% relaxation (F_0.5max_), time to minimum first derivative (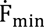), and time to peak contraction (F_max_). Points are color-coded per the scale at right, indicating the log_10_[TTX] IC_50_ of each individual as found in **a,e**. All p-values were calculated by Kruskal-Wallis nonparametric analysis of variance.

To test for a trade-off with increasing TTX resistance, we evaluated the force and speed of toxin-sensitive and toxin-resistant muscles grouped by species and genotype. To avoid potential genetic background-dependent effects, we only made relevant comparisons within species. The mean twitch force produced by skeletal muscles carrying TTX-resistant Na_V_1.4 was three-fold (*Th. atratus*) and five-fold (*Th. sirtalis*) less than TTX-sensitive skeletal muscles (Fig. 3b,f, Supp. Fig. 4c,h). The peak rate of force development (Ḟ_max_), which is known to be closely related to action potential generation and therefore sodium channel performance^67,69^, is reduced accordingly (Supp. Fig. 4b,g). Close attention to contraction timing generally shows that deficits in the speed of force development compound throughout the progression of any single twitch (Fig. 3i,j) with variable effects on overall contraction duration (Supp. Fig. 4d,i).

Differential effects of toxin resistance mutations on the progression of force development and contraction timing were observed in each snake species. For toxin-resistant *Th. atratus* (Fig. 3i) carrying Na_V_1.4^EPN^, muscles reached a premature maximum force that quickly resolved back to baseline levels (Fig. 3i, Supp. Fig. 4d,i). However, force development was definitively delayed in *Th. sirtalis* carrying Na_V_1.4^LVNV^ (Fig. 3j). These unique p-loop mutations exert distinct effects on contraction timing that scale up to more complex activation as well, such as in tetanus. While peak tetanic force was reduced in both resistant populations (Fig. 3c,g), only resistant *Th. sirtalis* (Na_V_1.4^LVNV^) failed to develop force as quickly as their TTX-sensitive counterparts (Supp. Fig. 4e,j). This slower rate of force generation is consistent with the electrophysiological finding that only Na_V_1.4^LVNV^ entered steady-state inactivation more quickly than ancestral channels (Supp. Fig. 1g,h). Presumably, as a tetanic contraction progresses, a larger proportion of the channel population inactivates, preventing sustained rapid force development. This likely gave rise to the slower morphology of the mean tetanic contraction in contrast to any of the other species-genotype comparisons.

Comparing peak force (F_max_) to TTX IC_50_ revealed a clear trade-off between resistance-conferring and ancestral genotypes at the muscular level (Fig. 3d,h). Greater toxin resistance in muscles is accompanied by reduced muscular force in both species, consistent with biophysically compromised channels. Therefore, the pore domain mutations in *Th. atratus* and *Th. sirtalis* possess both gain-of-function and loss-of-function features, although it remains unclear which individual substitutions contribute gains in toxin resistance or losses in excitability. As such, we conclude that mutations altering the structure of the sodium channel pore coincide with reduced channel function and concomitant tetrodotoxin resistance. Skeletal muscles bearing channels with these altered structures also show reduced performance despite extraordinary tetrodotoxin resistance. These results sufficiently explain previous findings that some toxin-resistant snakes are slower than their toxin-sensitive counterparts ^24^.

### Resistance mutations perturb the stability of the selectivity filter and S5/S6 helices

Molecular modeling of the sodium channel provides insights into mechanisms that likely contribute to the reduced unitary conductance of toxin-resistant channels. These biophysical findings may also explain why the quadruple-point mutant (*Th.s.*Na_V_1.4^LVNV^) enters into steady-state inactivation faster than either the toxin-sensitive (*Th.*spp.Na_V_1.4^+^) or triple-mutant toxin-resistant (*Th.a.*Na_V_1.4^EPN^) proteins. To understand the molecular stability of these three channel variants, the crystal structure of the human sodium channel was modified to create a homology model of the ancestral snake protein^17,70^. This ancestral version was then further modified to create homology models of both Na_V_1.4^EPN^ and Na_V_1.4^LVNV 13^. These three models were then run through an optimization routine. We compared the stability (ΔG) of key residues between these three protein models, including the resistance-conferring sites, the selectivity filter, the activation and inactivation gates, and hydrophobic residues at the interface of the pore-forming loops and the S5/S6 transmembrane passes, that cumulatively form the ion-conducting pore^71–73^.

Calculating the total interaction energy of residues in the ancestral channel and contrasting to mutant toxin-resistant channels demonstrates that the toxin-sensitive channel is the most stable protein, exhibiting a cumulative interaction energy of -393.85 kj/mol across all six residues (sites 1248, 1252, 1527, 1532, 1539, and 1540, using the human numbering system). While the triple-point mutant, Na_V_1.4^EPN^, is comparably stable at these sites (-366.84 kj/mol), the quadruple-point mutant, Na_V_1.4^LVNV^, is substantially destabilized (-42.43 kj/mol). In fact, Na_V_1.4^+^ and Na_V_1.4^EPN^ demonstrate residue pairings whose combined interaction energies are therefore 6.43x and 6.30x more stable, respectively, than the sum of the interaction energies between the same six sites in Na_V_1.4^LVNV^ (Supp. Table 12).

When decomposing these total interaction energies into their component interactions, it becomes clear that N1539 strongly destabilizes both Na_V_1.4^EPN^ and Na_V_1.4^LVNV^, accounting for the loss of 122.91 kj/mol stability in Na_V_1.4^EPN^ and 148.54 kj/mol in Na_V_1.4^LVNV^. However, in Na_V_1.4^EPN^, E1248 counteracts the instability conferred by N1539. While the conserved residues at positions 1527, 1532 and 1540 each have small, destabilizing interactions with N1539, E1248 interacts favorably with all four residues (I1527, I1532, N1539, and G1540). In fact, the most destabilizing interaction for E1248 is with its nearby neighbor, P1252, also found in resistant *Th. atratus* channels. So, while D1539N reduces the total favorable interaction energy at site 1539 by 86.4% in Na_V_1.4^EPN^, D1248E increases the favorable interaction energy by 353.6%. Therefore, E1248 appears to act as a compensating force against the destabilizing action of N1539.

In Na_V_1.4^LVNV^, however, no such countervailing force exists. While D1248 and N1539 appear to have a small, mutually stabilizing interaction (-8.58 kj/mol), the sum of interactions at sites 1527, 1532, 1539 and 1540 produce a net destabilizing effect. The majority of lost stability in Na_V_1.4^LVNV^ (ΔΔG=+351.41 kj/mol) can be explained by destabilizing interactions between three hydrophobic pairings (L1527-F412, V1532-Y1581, and V1540-G1512) and a hydrophilic interaction (N1539-K1244) (Figure 4). L1527 most likely interferes with the Domain I pore loop residue, F412, due to crowding by the bulky dimethyl group of leucine compared to the single methyl terminus of isoleucine. However, changes in residue volume may not completely explain why V1532 interacts negatively with the aromatic Y1581 of Domain IV S6. Rather, V1532 possesses a stronger attraction to a nearby L1541 in the ascending pore loop than isoleucine does in the toxin-sensitive channel. V1532 is consequently contorted by its interaction with L1541 and interferes with the hydrophilic hydroxyl group on Y1581, thereby yielding an unfavorable interaction. Further, V1540 negatively interacts with G1512 found on an extracellular loop, mostly likely due to steric hindrance compared to the ancestral glycine. Lastly, N1539 demonstrates a deleterious interaction with the positively charged residue, K1244, a key residue that constitutes the Domain III selectivity filter (the “K” of DEKA). This unfavorable interaction between N1539 and the Domain III selectivity filter is present and similar in magnitude in both Na_V_1.4^EPN^ and Na_V_1.4^LVNV^.

**Figure 4.**
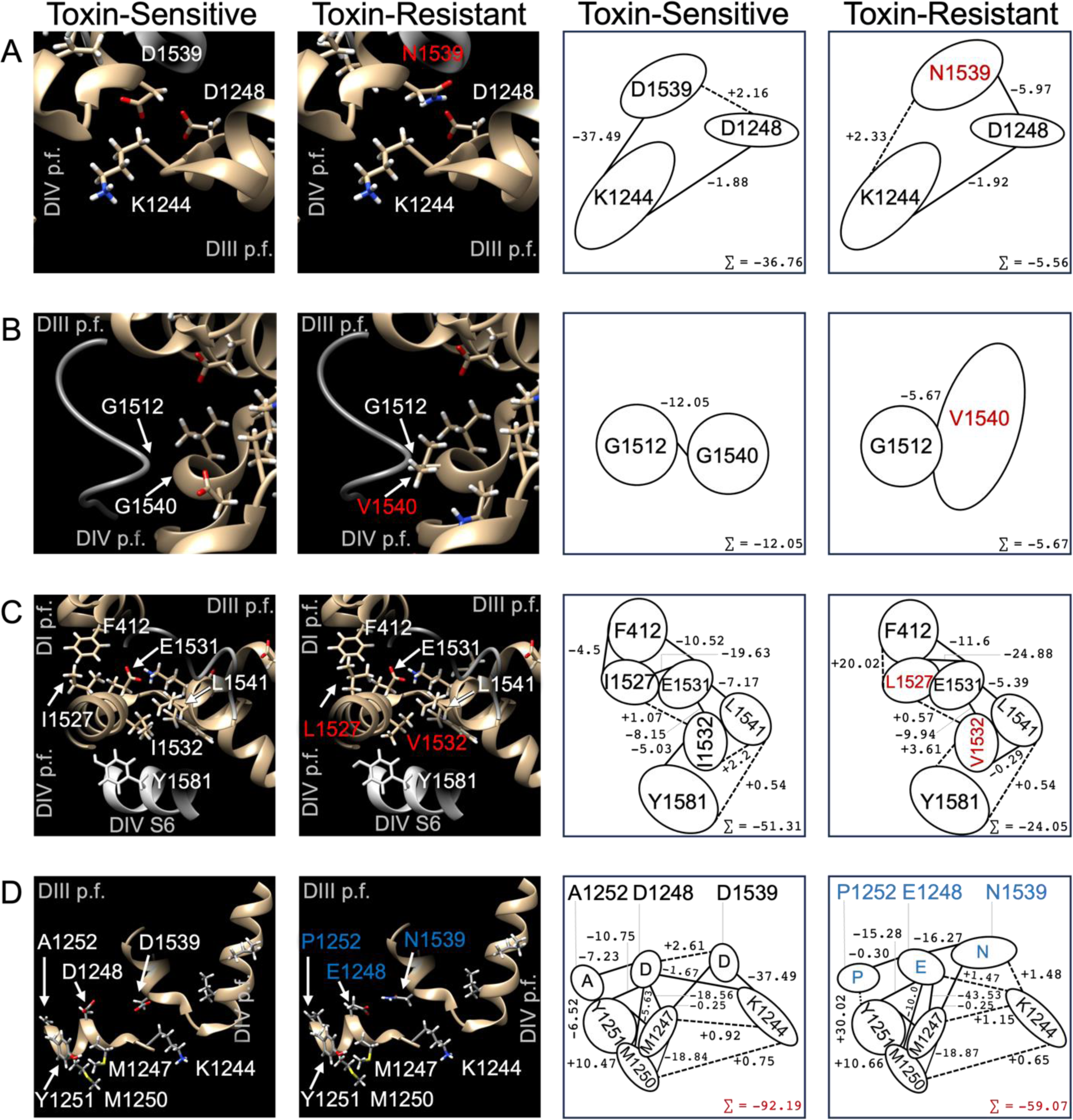
Network of interaction energies (ΔG, kj/mol) between key residues in sodium channels. Toxin-sensitive channels (left column) display different interaction energies compared to toxin-resistant channels (NaV1.4LVNV: rows A, B, & C; NaV1.4EPN row D). The left two columns show key scenes in the protein while the right two columns show simplified schematics demonstrating the interaction energies between residues of interest.

In addition to directly altering channel pore topology, mutations in TTX-resistant sodium channels appear to have indirect effects on channel function by destabilizing conserved residues nearby. Specifically, resistance mutations reduce the stability of the hyper-conserved DEKA residues that form the selectivity filter (D406, E761, K1244, and A1536)^39^. Though not directly mutated, the selectivity filters of the toxin-resistant channels are indeed less stable (-7.16% in Na_V_1.4^EPN^ and -10.7% in Na_V_1.4^LVNV^)(Supp. Table 13). The reduced stability of the ion-conducting pathway may contribute to the reduced conductance of the mutants without negatively affecting sodium selectivity ^43,64^. We found the toxin-sensitive channel’s median unitary conductance to be 15 pS, which corresponds to conducting around 9400 sodium ions per millisecond (at +10 mV). Presumably, this conductance is facilitated by a stable and rigid pore structure. Future molecular dynamics work will be needed to calculate the flow rate through a less stable pore. Such modeling may provide a full explanation of the reduced unitary conductance in toxin-resistant channels^74^.

Examining how the network of residue-residue interactions shifts from variant to variant may also help explain, at least from a structural perspective, why steady-state activation is unaffected by pore-domain mutations, and also why Na_V_1.4^LVNV^ does enter into steady-state inactivation faster than either Na_V_1.4^+^ or Na_V_1.4^EPN^. Pore-domain mutations on the extracellular face would not be expected to exert direct effects on the activation and inactivation gates near the intracellular face. Indirectly, however, the interior aspect of the pore loops interface with the S5 and S6 helices through hydrophobic interactions, which in turn, directly interact with the activation and inactivation gates (Figure 5). Thus, some aspects of steady-state kinetics can be mechanistically explained by interaction energies between the p-loops and the S5/S6 helices.

**Figure 5.**
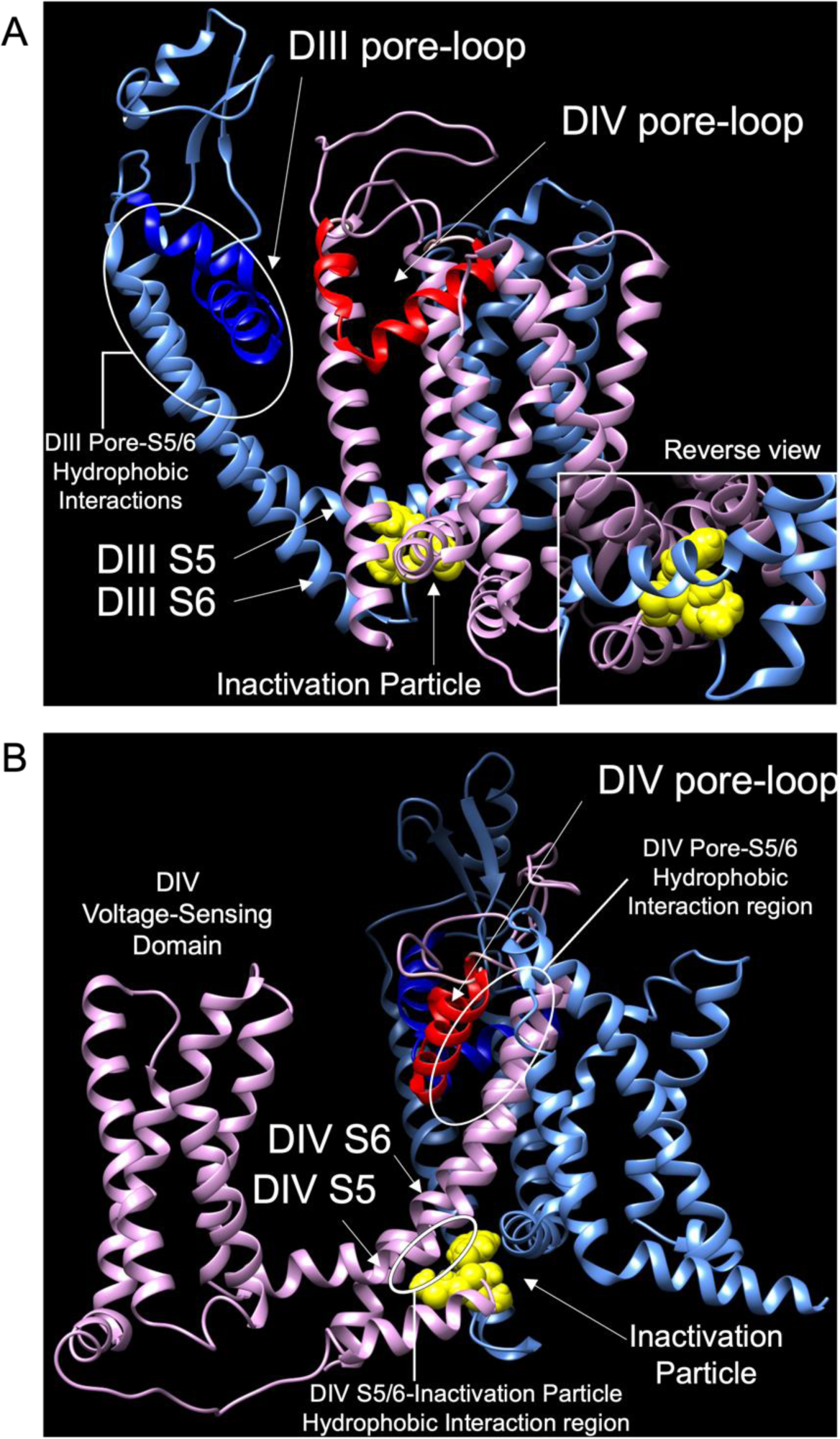
Scenes in the voltage-gated sodium channel depicting interfacing areas between the pore loops and the S5/S6 helices. The Domain III (A) pore loops (dark blue) interact with the S5/S6 helices (pale blue) via hydrophobic interactions from the interior aspect of the reentrant loops and the superior aspect of the S5/S6 helices. The insert shows how these same DIII S5/S6 helices interact with the inactivation particle (yellow). Similarly, the Domain IV (B) pore loops (red) interact with the DIV S5/S6 helices (pink), also through hydrophobic interactions. The DIV S5/S6 helices also directly interface with the inactivation particle. Therefore, the pore loops are indirectly linked to the inactivation gate via hydrophobic interactions.

The activation gate, a group of hydrophobic amino acids near the intracellular face of the pore, is dilated following the molecular reorganization of the voltage-sensing domains during depolarization. The activation gate in our modified Na_V_1.4 model is composed of Y450 (Domain I S6), F805 (Domain II S6), F1298 (Domain III S6), and F1601 (Domain IV S6)^75,76^. Our electrophysiological data show that activation is identical across all three channels. Likewise, the total interaction energy of the activation gate residues is identical in all three molecular models. Steady-state activation is therefore not negatively affected by pore-domain mutations.

Inactivation, however, appears to differ across these channels. Our electrophysiological and myographical data suggest that Na_V_1.4^LVNV^ inactivates prematurely. At rest, the inactivation gate (I1310, F1311, M1312) is locked in place by interactions with the S5 and S6 helices of Domains III and IV (Figure 5). From the intracellular face, the IFM particle interacts through hydrophobic forces with A1152 (DIII S5) and F1305 (DIII S6). From above, closer to the ion-conducting pathway, the IFM particle interacts, again through hydrophobic forces, with A1481 (DIV S5) and I1594 (DIV S6). As expected, with the pore mutations far away from the inactivation gate, the interaction energies between the IFM particle and the DIII and DIV S5/S6 residues are virtually identical across all channel variants studied here. Thus, differences in tendencies toward inactivation must arise from indirect effects of the pore-domain mutations, likely due to differences in the mobility of S5/S6 domains during the molecular reorganization that initiates inactivation.

The intracellular ends of the DIII and DIV S5 and S6 helices cradle the IFM particle while the extracellular ends of those same transmembrane α-helices form direct hydrophobic interactions with the interior aspect of the reentrant p-loops (Figure 6). For inactivation to initiate, the S5 and S6 helices must slide against one another to liberate the IFM particle, permitting it to contact and occlude the intracellular face of the pore^59^. Over both Domains III and IV, there are 44 key residues linking the p-loops to the S5/S6 helices. To mobilize the inactivation gate, Na_V_1.4^+^ channels must overcome the total interaction energy of those 44 residues (-4044.23 kj/mol) (Supp. Table 14). These residues in the triple-point mutant, Na_V_1.4^EPN^, are 76.92 kj/mol less stable than Na_V_1.4^+^, spread over both Domains III and IV. However, the p-loop-S5/S6 interactions are least stable in Na_V_1.4^LVNV^ (+109.06 kj/mol), whose mutations are all concentrated in Domain IV. This decreased interaction energy between the Na_V_1.4^LVNV^ pore-loops and the S5/S6 domains likely results in hypermobility of the DIV S5/S6 helices, thereby prematurely releasing the IFM particle to engage inactivation.

## Discussion

We demonstrate structural design limitations that constrain sodium channels from maximizing both biophysical performance and resistance to tetrodotoxin (TTX). The sites at play in adaptation to TTX serve a structural role in the sodium channel^77^, such that changes to these residues impose functional costs on the channel’s critical role in excitability. Accordingly, resistant *Thamnophis* skeletal muscles appear to bear the effects of poorly conducting channels in the form of reduced force and variable speed. Snakes rely on these sodium channels to maximize muscular and organismal performance, essential for foraging, wrestling with prey, escaping predators, hunting, and simply breathing. As such, key physiological and organismal tasks will be compromised by the reduced ionic conductance and myasthenic muscles observed in resistant snakes.

The reduced sodium channel unitary conductance reported here is novel, providing direct evidence of dramatic, traceable consequences to higher orders of organization. Though slow inactivation has been proposed as a biophysical penalty of toxin resistance mutations^43^, this functional cost is unlikely to underlie the scope and magnitude of skeletal muscle deficits we describe here. Furthermore, our physiological assessment of skeletal muscles in the snake-newt system provides a key link between molecular and organismal variation. Together with insights into the stability of key residues across the sodium channel variants studied here, our work ultimately provides a firm etiology of functional trade-offs in TTX resistant snakes that may ultimately limit phenotypic evolution.

The voltage-gated sodium channels of garter snakes exhibit similar but distinct evolutionary solutions and drawbacks to the steep physiological challenge of dietary TTX. Each solution is forced to preserve the critical function of the sodium channels on which organismal performance and survival depends. Both TTX resistance and muscular performance directly rely on channel pore structure, such that minute topological changes underlie broad phenotypic variation in an apparent zero-sum fashion. The biophysical insights from modeling the interaction energy of three channel variants demonstrates important constraints on protein structure and function, highlighting that even seemingly benign, conservative mutations can have cascading consequences throughout the protein.

The function of the sodium channel depends chiefly on the movement and relative spatial configuration of hydrophobic residues. The mutations seen in these snakes (Na_V_1.4^EPN^ and Na_V_1.4^LVNV^) appear to negatively impact function by interfering with the structural stability of the protein, resulting in hypermobility of these important hydrophobic residues in the pore and, in one case, a premature molecular reorganization that stops current flow. These molecular limitations on the types of mutations that are tolerable to protein function scale up to produce cellular and organ-level phenotypes that are likely suboptimal in many natural landscapes. Though other sodium channel mutations might offer perhaps even greater toxin resistance than Na_V_1.4^EPN^ or Na_V_1.4^LVNV^, such theoretical mutations likely breach the threshold of tolerable trade-offs to persist in nature. Resistance-conferring mutations in *Thamnophis* persist in just a narrow region of the sodium channel^23^ compared to the distribution of mutations in TTX-bearing species that must also be resistant to their own defenses^35–38^. In tetrodotoxic taxa, the persistent exposure to their own poison may require the incorporation of even more costly mutations. Garter snakes, on the other hand, may prey on newts irregularly ^78^, and as active, widely foraging, semi-aquatic predators^79^, may demand greater muscular performance, which likely acts as a brake on selection towards extreme TTX resistance.

Regardless, for these extremely resistant snakes, possessing such compromised proteins and impaired physiology may pose dramatic ecological and evolutionary consequences. Resistant snake performance may not meet the demands imposed by predators and prey in some ecological communities, or of harsh abiotic conditions, thereby limiting the geographic scope of resistance-conferring alleles and success of TTX resistance phenotypes. Snakes carrying *Th.a.*Na_V_1.4^EPN^ and *Th.s.*Na_V_1.4^LVNV^ may thus be confined to only those ecological communities or abiotic environments that can support the costly phenotypes those genotypes produce. Indeed, these TTX-resistant sodium channel genotypes are highly localized and only appear in certain “hotspots” rather than broadly distributed across the landscape as are toxin-sensitive animals^18,20,21,46,80^.

Antagonistic pleiotropy is thought to drive the convergent molecular evolution of TTX-resistant sodium channels^23,25^. Here, both snake species (*Th. atratus* and *Th. sirtalis*) have evolved the identical replacement D1568N ^13^, despite being distantly related^81^. Given that all proteins are universally subject to the chemical and physical forces that govern folding and function^5^, such biophysical constraints may ultimately underlie the degree to which evolutionary trajectories are predictable across lineages. Consequently, we conclude that the two suites of mutations (Na_V_1.4^EPN^ and Na_V_1.4^LVNV^) that have independently arisen in *Th. atratus* and *Th. sirtalis*, respectively, are the outcome of selection constrained by antagonistic pleiotropy to balance two opposing selective demands.

The trade-off we describe suggests that the structure-function relationship of the sodium channel is such that many possible mutations can reduce TTX blockade, but relatively few can do so while simultaneously maintaining sufficient levels of sodium channel function. This in turn must limit the number of mutations that satisfy two opposing selective demands imposed on garter snakes engaged in coevolutionary arms-races with newts. Such antagonistic pleiotropy therefore constitutes a limitation on adaptive evolution that, in the face of natural selection, promotes novel genetic variation in loci otherwise highly constrained by physical forces acting on the encoded proteins. Ultimately, simple biophysical limitations can manifest across levels of organization to shape molecular and phenotypic evolution in natural systems, potentially impacting the fates of populations and even species interactions.

## Materials and Methods

### Animal sampling and captive care

To measure whole-animal TTX resistance, as well as the properties of skeletal muscles in snakes with different sodium channel genotypes, we field collected individual *Thamnophis atratus* carrying Na_V_1.4^+^ (*Th.a.*Na_V_1.4^+^) and *Th. sirtalis* carrying Na_V_1.4^+^ (*Th.s.*Na_V_1.4^+^) for the ancestral state and their three- and four-point mutant counterparts (*Th.a.*Na_V_1.4^EPN^ and *Th.s.*Na_V_1.4^LVNV^). Our final sample included 62 animals from natural populations in California and Oregon, as well as a few lab-born individuals: 19 TTX-sensitive *Th. atratus* from three sites; 21 TTX-sensitive *Th. sirtalis* from six sites; 10 TTX-resistant *Th. atratus* from two sites; and 12 TTX-resistant *Th. sirtalis* were caught from four sites (Supp. Table. 1).

Field-caught *Thamnophis* spp. were kept in captivity at either Utah State University (USU) or University of Nevada, Reno (UNR) prior to any work. Snakes were provided water *ad libitum* and fed either previously frozen fish (tilapia or trout) or mice once or twice weekly with no detectable difference due to diet (data not shown). The snakes were kept at 26 ± 1 °C and on a 12L:12D photoperiod with hide boxes provided. All snakes were housed individually in mouse-style containers or aquaria commensurate with animal size. All cages had one end situated above heat tape to achieve a thermal gradient. All animal care procedures followed USU and UNR IACUC protocols (USU IACUC #1008 and UNR IACUC #00687).

### Organismal resistance assay

Animals were encouraged to sprint along a linear racetrack lined with artificial turf. Infrared sensors (every 20 to 50 cm) recorded snake progression along the 4 m track^14,82,83^. The two shortest time intervals between sensors were averaged and used to calculate the animal’s maximum speed. Snakes that refused to sprint down the track were omitted. All trials were performed at 26 ± 1 °C. Only two individuals (E.D.B., Jr. and C.R.F.) conducted sprint trials to reduce interrater variation.

This assay was repeated following intraperitoneal injection with TTX (MilliporeSigma, Burlington, MA, USA; Tocris, Bristol, UK; Alomone Labs, Jerusalem, Israel; dissolved in citrate buffer pH 6.8; injection volume ≤0.5 mL). All doses were calculated and delivered in mouse units. A mouse unit is the amount of TTX (2.857×10^-7^ g) required to kill a 20-g mouse in 10 minutes^84^. Scaling to account for total body mass of the snake to be injected, this mass-adjusted mouse unit (MAMU) contextualizes the snakes’ toxin resistance relative to TTX-sensitive mammalian tissue. Doses are calculated by the equation Dose (MAMU) = (*Dose* g TTX / *m* (g)) • (20 g / 2.857×10^-7^ g TTX), where *m* is the mass of the individual snake injected. A distribution period of 30 minutes was allowed before racing the individual. Control injections of physiological saline have no effect on snake performance (Brodie and Brodie 1990), and TTX resistance is not affected by repeated short- or long-term exposure^83,85^. No snake received more than five (5) injections, and all doses were separated by 48 hours to ensure the elimination of residual TTX and to reduce the effect of fatigue^83^.

Animals were not given standard doses of TTX but rather dosing was informed by the population of origin. We injected TTX starting at 1 MAMU for snakes from known low-resistance populations and 10 MAMU from known high-resistance populations. The dose was then serially increased (3, 5, 10 and sometimes 15 or 25 MAMUs for low-resistance; 25, 50, 100 or 250 MAMUs for known high-resistance). For individuals with exceedingly great resistance, the dose-response protocol was truncated to 250 MAMU to avoid potentially irreversible damage to the animal and preserve costly toxin reserves. The dose of TTX needed to slow an individual to 50% of its baseline sprint speed is reported as the 50% MAMU (analogous to a 50% inhibitory concentration, IC_50_).

Here, 50% MAMU was calculated by fitting the dose-responses to a sigmoidal curve. A custom R script was designed to handle this fitting and used the equation *Speed (m/s)* = A_2_ + (A_1_-A_2_)/(1+e((log(*Dose (MAMU)*)-log(x_0_))/dx)), where A_1_ and A_2_ are the upper- and lower-bounds, x_0_ is the center of the curve (50% MAMU) and dx is the rate of decline with increasing dose. The dose for baseline speed (0 MAMU TTX) was revised to 0.1 MAMU to abide by the limitations of the natural logarithm. A minimum of five (5) dose responses is needed to fulfill this model. Where conditions precluded the administration of more than three (3) doses of TTX, the sigmoidal fit was forced by including sufficiently large doses where speed is assumed to converge to zero m/s.

### Genotyping

Regions of *SCN4A* corresponding to the third and fourth pore domains of Na_V_1.4 (DIII-PD & DIV-PD) were sequenced for all snakes. Genomic DNA was extracted from tail snips, skeletal muscle, or liver tissue by DNeasy Blood & Tissue Kits (Qiagen Inc., Germantown, MD, USA). Fragments of genomic DNA corresponding to DIII-PD & DIV-PD of *SCN4A* were amplified by PCR using *Thamnophis*-specific primers (Supp. Table 2)^13,20^. Fragments were submitted to the Nevada Genomics Center (UNR) or the DNA Analysis Facility (Yale University) for bidirectional Sanger sequencing using amplification primers, and sequences were resolved on an Applied Biosystems Prism 3730 DNA Analyzer or 3730*xl* DNA Genetic Analyzer. Nucleotide sequences were aligned and translated to amino acid sequences using a variety of software, including Geneious versions 4.8.5 and 6 (Biomatters Ltd., Auckland, New Zealand; http://www.geneious.com) or Sequencher 4.2 (Gene Codes Corp., Ann Arbor, MI, USA), ClustalW 1.83 (Clustal W and Clustal X version 2.0) and MacClade 4.08^86–88^. We deposited all sequences in GenBank (accession numbers: Exon 22 ON609732-ON609763; Exon 26 ON609764-ON609824).

Animals from both *Th. atratus* and *Th. sirtalis* were scored as Na_V_1.4^+^ (TTX-sensitive) for translated *SCN4A* sequences reading 1276-MDIMYA-1281 in DIII and 1556-ICLFEITTSAGWDG-1569 in DIV. *Thamnophis atratus* were scored Na_V_1.4^EPN^ (TTX-resistant) when exhibiting D1277E and A1281P in DIII and D1568N in DIV. *Thamnophis sirtalis* were scored Na_V_1.4^LVNV^ (TTX-resistant) when exhibiting I1556L, I1561V, D1568N, and G1569V. Snakes confirmed as heterozygous with respect to *SCN4A* were omitted from further testing.

## Dissection

Immediately prior to myography work, animals were humanely euthanized without the use of chemicals that could affect sodium channel performance. Euthanasia was performed independent of circadian rhythm and without attention to feeding schedules or shedding cycles. All animal procedures followed UNR IACUC protocols (IACUC # 00687). The specimen was submerged at all times in fresh Krebs buffer and supplied with 95%O_2_/5%CO_2_. A segment of 2 to 4 cm of the snake body was transferred to a dish supplied with the same solution and gasses for dissection under an upright stereomicroscope. Cleaved down the ventral midline, the segment was skinned and pinned to the dish by the ribs and intercostal muscles. The ventrolateral muscle group known as the *iliocostalis* muscle was extracted using curved-blade microdissection scissors. All manipulations of the muscle were made to minimize trauma and any undue tension on the muscle prior to suspension, typically in under three minutes.

### Skeletal muscle performance and resistance assay

Myography was performed on a Contraction System Chamber 800A which was operated by Dynamic Muscle Control DMC v4 software (Aurora Scientific Inc., Aurora, Ontario, Canada). Tensile strength was recorded by an Aurora Scientific 300C-LR Dual Mode Lever System. Before suspending the *iliocostalis* muscle, the force transducer was calibrated using a 40 g load. A 0.15 mm nylon suture was driven through the muscle group and secured by modified Timber Hitch knots. The chamber was prefilled with Krebs buffer, continuously perfused with 95%O_2_/5%CO_2_, and maintained at 25 °C by water-jacketed circulation (Lauda Alpha A6, Germany). The buffer was allowed to equilibrate for five minutes prior to submerging the muscle. Electric field stimulation (EFS) platinum electrodes ran parallel along the length of the muscle which was situated equidistant from both electrodes.

A baseline tension of 0.1 g was applied to the suspended muscle. The EFS was optimized by applying a 500 μs pulse of direct current in increasing magnitude from 1 to 300 mA. The EFS stimulus was set to 1.5x the smallest magnitude current that provided the greatest recruitment of the muscle. At this optimum electrical stimulus, the length-tension relationship was optimized by increasing the baseline tension until the force output peaked. A 1 min rest period was allowed between the end of the optimization routine and the beginning of the performance trials. All trials were recorded at 10 kHz.

#### Transient Muscular Force Assessment

Four consecutive EFS pulses, one every five seconds, were applied to the muscle under the optimized conditions as described above. These traces are the basis of comparison for muscular force generation between genotypes and TTX resistance phenotypes within *Thamnophis* spp. The resulting transient contractions were analyzed for peak force magnitude (N/g of tissue), contraction duration (from center of the stimulus to 50% relaxation), and latency (time from center of the stimulus to the peak force generated). The first derivative of each trace was analyzed for its peak rates of force development and relaxation (kN/g s^-1^). Of the train of four pulses, comparisons were only made between contractions of the same order.

#### Tetanic Muscular Force Assessment

Under the same optimized EFS conditions as the transient force assay, the muscle was stimulated at 1000 Hz for 2 seconds. The resulting tetanic contraction was analyzed for peak force magnitude (N/g of tissue), contraction duration (from stimulus onset to 50% relaxation), and latency (time from stimulus onset to the peak force generated). The first derivative of each trace was analyzed for its peak rates of force development and relaxation (kN/g s^-1^). The tetanus routine is so stressful that it is terminal. A new muscle was dissected and used following this protocol. Transient force, rheobase, and tetanic force protocols were run within a period of 15 minutes with 1 minute of rest between protocols.

#### Skeletal muscle TTX resistance assay

The transient muscular force assessment and EFS optimization were repeated with a new, freshly dissected muscle. The optimal transient was stimulated and recorded. Known quantities of TTX dissolved in citrate buffer pH 6.8 were added to a bath of known volume. The doses began as low as 1 nM and increased until the transient force peak magnitude decayed. In many cases, [TTX] was increased until the absolute abolishment of muscular activity was achieved, though this was not possible for every experiment. The volume of TTX in citrate buffer added to the bath was kept below 0.01% of the total bath volume to minimize the effects of diluting the Krebs buffer. However, in cases when exceedingly great [TTX] was needed to inhibit the contraction and stock concentrations of TTX dilutions were too low to keep the volume added below 0.01% of the bath volume, a separate experiment where an identical volume of citrate buffer (without TTX) was added to verify the absence of dilution-related effects.

Dose response curves were generated from the peak transient contraction force magnitude. The responses were fit to a sigmoidal curve of the equation *Force (N/g)* = A_2_ + (A_1_-A_2_)/(1+e((log(*[TTX] (nM)*)- log(x_0_))/dx)), where A_1_ and A_2_ are the upper- and lower-bounds, x_0_ is the center of the curve (toxin concentration at half-maximal inhibition, IC_50_) and dx is the rate of decline in force with increasing TTX concentration. Comparisons of IC_50_ (nM) and dx (N/g nM^-1^) were restricted to animals within the same species, grouping animals by genotype. To abide by the limitations of the natural logarithm, the dose at baseline transient force magnitude (0 nM TTX) was revised to 0.1 nM. If the contraction recorded at 0 nM TTX was unusable or unavailable, the dose responses were normalized to the maximum force produced throughout the routine (frequently occurring within the first 10 nM TTX). Dose response data could not be fit with fewer than four concentration-force points collected. However, when only three points were available, a fourth point was fabricated where the force would visually converge to zero N/g (*i.e.,* 100 µM, which abolishes contractility in even the most resistant snake muscles).

### Blinding and randomization

No animals had confirmed genotypes at the time of the whole-animal performance and resistance assays. For *Th. atratus*, all animals were processed blindly and genotyped post hoc to prevent rater bias. For *Th. sirtalis*, genotypes were scored prior to the experiment, and an investigator (JSR) made a randomized and blinded list of snakes to process. This list was handed off to the investigator performing the myography experiments (REdC) unaware of any animal’s genotype and whole-animal phenotype until all animals on the list were processed.

### Assessment of Na_V_1.4 function across genotypes

#### Mammalian Cell Culture

Adherent Human Embryonic Kidney 293 cells (HEK293, authenticated from vendor, ATCC® CRL1573^TM^, Manassas, VA, USA) were maintained in Dulbecco’s modified Eagle’s medium (DMEM, ThermoFisher Scientific, Waltham, MA, USA) supplemented with 10% v/v heat-inactivated fetal bovine serum (Atlanta Biologicals, Flowery Branch, GA, USA) and 100 U/mL penicillin/streptomycin (ThermoFisher Scientific, Waltham, MA, USA). The cells tested negative for mycoplasma (validated by ATCC and by the authors using Hoechst 33258 staining). Cells were not assessed for cross-contamination by other mammalian cell lines, as our electrophysiological methods are robust to any consequent differences. Cells were plated on nonpyrogenic culture vessels (Sarstedt, Nümbrecht, Germany) and incubated at 37°C supplied with 13.0% CO_2_ for pH 7.3 at 100% humidity. The age of the cell populations ranged from approximately 7 to 20 passages. All passages were carried out with TrypLE Express trypsin reagent (ThermoFisher Scientific, Waltham, MA, USA). HEK293 is a commonly misidentified cell line (ICLAC) but was used here only as a representative eukaryotic expression system. Cells were only used for transient lipofection and expression of sodium ion channels and subsequent patch clamp electrophysiology.

#### Plasmid construction

An oocyte-expression plasmid containing a *Rattus norvegicus* sodium channel clone (*rSCN4A*) was kindly provided by Mohammed Chahine (Université Laval, Québec, Canada). A rat clone was deemed acceptable for preliminary research provided that the pore domains are entirely conserved between rats and snakes. The insert was extracted by EcoRI restriction enzyme digestion and inserted into a bicistronic mammalian expression vector, pIRES-hrGFP-2a (Agilent, Santa Clara Co., CA, USA). The new vector was then transformed into 5-alpha competent high-efficiency *E. coli* (strain C2987I, New England Biolabs, Ipswich, MA, USA) and amplified. The pIRES-hrGFP-2a-r*SCN4A* plasmid was then isolated by mini-prep (Qiagen Inc., Germantown, MD, USA).

#### Site-directed mutagenesis

The mutations made to pIRES-hrGFP-2a-r*SCN4A* are identical in relative position and amino acid substitution to those in *Thamnophis SCN4A* and in some cases converge with tetrodotoxic *Taricha* and many tetrodotoxin-defended fishes of the Tetraodontidae. We specifically focused on *SCN4A* mutations and excluded peripheral nerve-type sodium channels (*SCN8A*/Na_V_1.6 and *SCN9A*/Na_V_1.7) because all garter snakes carry the same mutations in those channels^24,89^.

The Na_V_1.4^EPN^ channel variant naturally occurring in *Th. atratus* was generated by site-directed mutagenesis (QuikChange Lightning II, Agilent, Santa Clara Co., CA, USA; now Stratagene, La Jolla, CA, USA). Using Agilent’s online primer design tool, primers were designed to implement the following changes: D1277E and A1281P in DIII and D1568N in DIV. In the pIRES-hrGFP-2a-*SCN4A* clone, these amino acid substitutions were achieved by t4640g, g4650c, and g5511a^18^. Mutagenic primers were synthesized, and their sequences can be found in Supp. Table 2.

Another variant was generated by *Mutagenex* (Suwanee, GA, USA) to include the mutations found in TTX-resistant *Th. sirtalis*: Na_V_1.4^LVNV^, I1556L, I1561V, D1568N, and G1569V in DIV^17^. In the pIRES-hrGFP-2a-*SCN4A* clone, these amino acid substitutions were achieved by a5475c, a5490g, g5511a, and g5515t. Mutagenic primer designs for this mutant were not reported, but the final sequence was confirmed. All three plasmid preps were then amplified by the same service (plasmid.com, Fargo, ND, USA) to obtain comparable DNA quality. Mutagenesis verification for all constructs has been submitted to GenBank (access: ON609825-ON609830).

#### Transfection

Cells were transiently transfected pursuant to the Lipofectamine 3000 protocol (ThermoFisher Scientific, Waltham, MA, USA). HEK293 cells were passed onto a 35 mm dish 24 h in advance to be 70% confluent at the time of transfection. Cells were transfected with 1 µg of pIRES-hrGFP-2a-r*SCN4A* or its mutants and allowed to translate protein for 48 to 72 h. This protocol was optimized on the wild-type clone and produced indistinguishable fluorescence reporter activity across all variants tested. Transfections took place within a 4-week period, and the plasmid variants were transfected in random order throughout the experimental period to avoid time-related effects on expression.

#### Electrophysiology

Clampex of the pCLAMP 10.7 software suite and an Axopatch 200A amplifier with CV202A headstage (Molecular Devices, Sunnyvale, CA, USA) were used to design and carry out voltage-clamp protocols and record *I_Na_.* An online 10 kHz Bessel filter was applied to the signal, and the resultant *I_Na_* was digitized at 50 kHz (Axon Digidata 1440A, Molecular Devices, Sunnyvale, CA, USA). Pipettes were fashioned from (1.5 mm OD, 1.1 mm ID) borosilicate capillary glass (BF150-110-7.5, Sutter Instruments Company, Novato, CA, USA). All patch pipettes and sharp electrodes were pulled on a P-87 Flaming/Brown Micropipette Puller (Sutter Instruments), and patch pipettes were polished on a Narishige MF-830 microforge. The pipette electrode resistance varied from 2-5 MΩ with 2.8 MΩ median resistance. The range of cell capacitance was measured to be between 5 and 14 pF. Series resistance was compensated to the edge of oscillation, and tau was set to 3 μs for all cells.

Cells were enzymatically dissociated and plated at 10% confluency. Successfully transfected single HEK293 cells were identified by green fluorescence during irradiation with 488 nm light filtered from a mercury lamp. Prior to making membrane contact, a correction was made for pipette capacitance and a liquid junction potential between internal and external solutions (-4 mV). The whole-cell patch clamp configuration was used to record macroscopic Na^+^ currents (*I_Na_*). Following the attainment of a gigaOhm seal, a five-minute rest period was allowed for dialysis of the internal solution and further stabilization. The ambient temperature was 23 ± 2 °C. Series resistance was compensated to ≥90%, τ = 1-5 μs.

The membrane surface area (pF) was estimated by integrating the capacitive current generated by a brief +10 mV pulse. The voltage dependence of *I_Na_* (I-V) was assessed by 10 ms test pulses from -80 to +80 mV in 10 mV increments. The test pulses followed a 500 ms conditioning period at -100 mV to remove inactivation. Following the test pulse, the membrane potential was returned to -100 mV to provide a symmetric but opposite capacitive current for reference. Currents were normalized to the membrane surface area to provide current density (pA/pF).

#### Channel kinetics

Steady-state activation (SSA) curves were constructed from the voltage dependence of G_Na_. This follows G_Na_= *I_Na,max_*/ (E_m_ – E_rev_), where E is the membrane potential during the test pulse and E_rev_ is the empirical reversal potential. Steady-state inactivation (SSI) curves were constructed by plotting *I_Na,max_*(pA/pF) against each prepulse potential. The resulting points were fit to the Boltzmann equation G/G_Na,max_ or *I_Na,max_* = 1/[1+exp(E_1/2_-E_m_)/k_v_], where E_1/2_ is the voltage that half-maximally activates or inactivates an Na_V_1.4 population and k_v_ is the slope factor. To pool curves from multiple cells, these curves were normalized to the upper bound and forced to approach 0% (for SSI) and 100% (SSA). Window currents are a byproduct of overlapping SSA and SSI behaviors whereby small populations of sodium channels tend to linger in the open state. Differences here could be indicative of hazardous biophysical trade-offs. The E_1/2_ and k_v_ parameters from the SSA and SSI curves can predict the Na_V_1.4 open probability during the window period. This is done using the equation P_open,window_ = [1/(1+e((E_1/2,SSA_ – E_m_)/k_v,SSA_)) • [1/(1+e((E_m_-E_1/2,SSI_)/k_v,SSI_)).

#### Assessment of recovery from inactivation

A paired-pulse protocol was applied to the membrane to assess the time and voltage dependence of recovery from inactivation. The population was first equilibrated (500 ms conditioning prepulse at -100, - 80, or -60 mV) and activated (10 ms test pulse to +10 mV, at peak SSA for all mutants), thereby engaging inactivation. The membrane was then returned to the prepulse potential before a second test pulse (10 ms, +10 mV), with the duration between test pulses serially increasing until the peak current was completely rescued. The ratio (*I_Na,max_*)_test-2_/(*I_Na,max_*)_test-1_ was plotted against the interpulse interval. The time constant of the recovery from inactivation (τ*_rfi_*) was calculated by a single exponential fit (OriginPro 2017). Comparisons were made between Na_V_1.4^+^, Na_V_1.4^EPN^, and Na_V_1.4^LVNV^ for each prepulse potential.

#### Onset of fast inactivation

The membrane was conditioned at -100 mV for 500 ms to remove inactivation before each sweep of this protocol. An initial prepulse potential at -30 mV for 0.2 ms preceded a test pulse at 0 mV for 10 ms. All alleles of Na_V_1.4 have been shown to completely inactivate for sufficiently long exposures to this depolarized potential. The time course of the onset of this inactivation was assessed by repeating this command 100 times, each time increasing the prepulse potential by 0.5 ms. The *I_Na,max_* of the current developed following the 0.2 ms pulse at -30 mV was taken as the maximum. The maxima of all sweeps were reported as a fraction of this first peak. The fraction of available channels was then plotted against the duration of the inactivating pulse. The time constant of the onset of fast inactivation (τ*_ofi_*) was calculated from a single exponential function and is the basis of comparison between alleles.

#### Fluctuation Analysis

The cell membrane was conditioned at -100 mV for 20 ms prior to a brief 10 ms pulse to +10 mV. Na_V_1.4^+^, Na_V_1.4^EPN^, and Na_V_1.4^LVNV^ are known to have saturated steady-state activation at this test voltage. This protocol was repeated 1000 times, and the subsequent mean current (*I_Na_*(t), pA/pF) was plotted against the variance over time (σ*_INa_*(t)^2^, pA/pF^2^). The data were fit by a parabola of the equation: σ*_INa_*(t)^2^ =[*I_Na_*(t)]i – [*I_Na_*(t)]^2^/N, where t spans from the time *I_Na,max_* occurs to the end of the test pulse (∼8.5-9.5 ms), i refers to the predicted unitary current of Na_V_1.4 and N refers to the predicted number of such channels in the tested membrane.

#### Resistance of Na_V_1.4 Alleles

The cell membrane was conditioned at -100 mV for 500 ms prior to a 10 ms test pulse to +10 mV. TTX was applied by perfusing aliquots of extracellular solution to which known volumes of TTX diluted in citrate buffer pH 6.8 were added. Added volumes were kept below 0.01% of the aliquot volume. The cell was allowed to equilibrate for two (2) minutes at the intended [TTX] (in nM: 0, 100, 200). Each current trace was then normalized to *I_Na,max_* (pA/pF) of the 0 nM trace.

#### Data analysis

Currents were analyzed with pCLAMP software 10.7 (Molecular Devices, Sunnyvale, CA, USA). Important metrics were recorded in Microsoft Excel (Redmond, WA, USA). Statistical tests and tests for approximate normality were performed in R (R Core Team 2017). Graphs were constructed in OriginPro 2017 (OriginLab Corp, Northampton, MA, USA).

#### Sodium Channel Molecular Modeling and Interaction Energy Estimation

To offer molecular insights into the differences between skeletal muscle voltage-gated sodium channels with different pore-domain residues, we turned to published crystal structures. The structure of the human voltage-gated sodium channel Na_V_1.4 in complex with β1 was used as a starting point (pdb 6agf, UniProt P35499). The sequence of the α subunit was then aligned (Blastp) to a consensus sequence of *Thamnophis* spp. skeletal muscle sodium channels from toxin-sensitive *Thamnophis sirtalis*, *Th. atratus*, and *Th. elegans*. The alignment revealed 356 differences between the *Homo* and *Thamnophis* sodium channels. In UCSF Chimera, all 356 point mutations were made to the human structure to generate a homology model that approximates the snake channel structure (ignoring differences in the C and N termini)^90^. This “serpentized” homology model therefore uses the human residue positioning numbers. Therefore, to recreate the mutations found in TTX-resistant snake Na_V_1.4 Domains III and IV of *Th. atratus* (D1277E, A1281P, and D1568N), the toxin-sensitive model was mutated at D1248E, A1252P, and D1539N. To recreate the Domain IV mutations of *Th. sirtalis* (I1556L, I1561V, D1568N, and G1569V), the toxin-sensitive model was mutated at I1527L, I1532V, D1539N, and G1540V. All three of these models were then run through UCSF Chimera’s standard Minimize Structure routine to produce three optimized, *Thamnophis* homology models for and Na_V_1.4^+^, Na_V_1.4^EPN^, and Na_V_1.4^LVNV^. Each of these models was then uploaded to the Interaction Energy Matrix (IEM) Web Application (https://ip-78-128-251-188.flt.cloud.muni.cz/energy/)^71–73^. For all six positions in question (residues 1248, 1252, 1527, 1532, 1539, and 1540), the interaction energy of each mutation was calculated by the IEM program for each minimized model, accounting for both Coulombic and Lennard-Jones modeled Van der Waals forces. Furthermore, the total interaction energies of several key residues linking the pore-domain and the S5/S6 domains of the sodium channel, that in turn link to the inactivation gate, were also calculated. The ΔΔG was calculated here using the ancestral, toxin-sensitive model as the reference. Residues that carry a more negative interaction energy (i.e. more stable) than the ancestral residue produce negative values while substitutions that destabilize the interaction energy relative to the ancestral protein produce positive values. Here, key residues were identified due to their known functional significance or by physical proximity when two atoms on adjacent residues lie within 5Å of one another.

### Solutions

All experiments on skeletal muscle tissue were performed in Krebs buffer of the following composition (in mM): 145 Na^+^, 127.7 Cl^⁻^, 25 HCO_3_^⁻^, 5.5 glucose, 5.4 K^+^, 1.2 Mg^2+^, 1.8 Ca^2+^, 1.2 H_2_PO_4_^⁻^, 1.2 SO ^2⁻^ and supplemented with 95% O_2_ and 5% CO_2_ to achieve pH ∼7.35. For whole-cell patch clamp recordings of HEK293 cells transfected with pIRES-hrGFP-2a-*SCN4A*, the pipette solution contained (in mM): 10 Na^+^, ≥120 Cs^+^, 110 aspartate^-^, 32 Cl^⁻^, 12 tetraethylammonium^+^, 5 EGTA^4⁻^, 5 HEPES, 3 Mg^2+^, and 1.5 ATP^4⁻^, adjusted to pH 7.2 with CsOH. The extracellular solution was formulated (in mM): 145 Na^+^, 160.6 Cl^⁻^, 10 tetraethylammonium^+^, 5.5 glucose, 5 HEPES, 1.8 Ca^2+^, and 1 Mg^2+^, adjusted to pH 7.4 with CsOH (final [Cs^+^] ∼ 1.6 mM). The measured liquid junction potential between the external and internal patch solutions was -4.4 mV (n = 18, data not presented), for which the pipette offset was adjusted to +4.4 mV prior to cell contact.

### Statistics

Mean data are presented alongside the standard error of the mean (sem). However, all statistical tests were performed using nonparametric analysis of variance in R 3.5.1 (R Core Team 2017) using Kruskal-Wallis ANOVA and Dunn’s multiple comparison post hoc test. Nonparametric methods were employed for all tests given that not all data were normally distributed. Statistical significance was determined by comparing the control and mutant groups using the Kruskal-Wallis test with α = 0.05. Preliminary experiments used to secure funding suggested that the effect size between resistant and sensitive animals would be sufficiently large to forego an effect size calculation or other power analysis. All test statistics and observation counts (n) can be found in Supplemental Tables 3-11 in the data repository below. All *p* values presented were adjusted to account for multiple tests by the conservative Bonferroni correction method. Center lines on box-and-whisker plots represent medians bounded by third and first quartiles; upper caps represent the 95^th^ percentile, while lower caps represent the 5^th^ percentile. Error bars represent the standard error.

## Data Availability Statement

Myography, whole-animal assays, and electrophysiology data that support the results can be found at https://github.com/rdelcarlo/Snake_Biophysical_Trade-off.git, in addition to tabular reports on statistical tests performed to form the conclusions presented here. All data sets and/or analyses generated in this study are available from the corresponding author upon reasonable request. All custom codes used to process these data can be found at the listed github.com.

## Author Contributions

C.R.F. conceived the hypothesis and is the principal advisor on evolution and ecology. N.L. is the principal advisor on biophysics, pharmacology, and physiology and, with R.E.d.C., designed the whole-cell recording protocols. C.R.F. and N.L. designed and performed the preliminary myography experiments that secured funding from the National Science Foundation (IOS-1355221). E.D.B.Jr., C.R.F. and H.A.M. carried out the whole-animal resistance assays, with essential aid from J.S.R. C.R.F., J.S.R., M.T.J.H. and E.D.B. III caught all wild animals, and C.R.F, M.T.J.H., J.S.R. and R.E.d.C. genotyped the animals used in this study. J.S.R. supervised the randomization and blinding of the myography experiments. H.A.M. carried out the animal husbandry. J.S.R. worked with R.E.d.C. to oversee the construction of a myography database and ensure data availability. S.A. directed the subcloning of the sodium channel gene into its plasmid vector. R.E.d.C. sequenced and mapped the sodium channel clone and generated or arranged the generation of site-directed mutants. R.E.d.C. carried out all myography and patch clamp experiments and performed all analyses and statistics. R.E.d.C. prepared all figures. R.E.d.C wrote the initial draft of this paper and reconciled all coauthors’ input into the final submission.

## Acknowledgments

We thank CAF&W for permits to C.R.F, M.T.J.H. and E.D.B. III, and we thank USU and UNR IACUCs for approval of live animal protocols. We thank Mike Edgehouse, Vicki Thill, Erica Ely, and Kevin Wiseman for aid with field collections, and Jeff Wilcox, John Bailey, Cathy Koehler, Pete Steel, and Margot Rawlins for access to field sites. We are grateful to Gabrielle Blaustein, Vicki Thill, Amber Durfee, Kenzie Wasley, Taylor Disbrow, Sage Kruleski and Aubrey Smith for aid in live animal care at UNR. We thank Mohammed Chahine for providing the wild-type rat Na_V_1.4 clone. Aspects of this work would not have been possible without the help of Christopher Lingle who provided key aid on non-stationary noise analysis. We thank Kimberly Stanek for producing crystal structure models of the sodium channel and Devon Picklum for snake photographs. Finally, we thank UNR Evol Doers for thoughtful discussions and useful feedback on this manuscript, especially Marjorie Matocq and Jamie Voyles. This work was supported by an NSF grant to C.R.F. and N.L. (IOS-1355221) and NIH grants to N.L. (R01 HL091238 and P20GM130459) and by an NIH grant to S.A. (R01 HL161122).

